# Tube2FEM: a general-purpose highly-automated pipeline for flow related processes in (embedded) tubular objects

**DOI:** 10.1101/2024.06.22.600203

**Authors:** Hani Cheikh Sleimana, Kevin M. Moerman, Diana C. de Oliveira, Joseph Jacob, Nesrin Mogulkoc, Brian R. Davidson, Simon Walker-Samuel, Rebecca J. Shipley

## Abstract

This paper presents a comprehensive and highly-automated open-source pipeline for simulating flow and flow-related processes in (embedded) tubular structures. Addressing a critical gap in computational fluid dynamics (CFD) and simulation sciences, it facilitates the transition from raw three-dimensional imaging, graph networks, or CAD models of tubular objects to refined, simulation-ready meshes. This transition, traditionally labor-intensive and challenging, is streamlined and highly-automated through a series of innovative steps that include surface mesh processing, centre-line construction, anisotropic mesh generation, and volumetric meshing, leading to Finite Element Method (FEM) simulations. The pipeline leverages a range of open-source software and libraries, notably *GIBBON*, *FEniCS*, and *Paraview*, to provide flexibility and broad applicability across different simulation scenarios, ranging from biomedical to industrial applications. We demonstrate the versatility of our approach through five distinct applications, including the mesh generation for soil-root systems, lung airways, microcirculation networks, and portal vein networks, each originating from a different data source. Moreover, for several of these cases, we incorporate Computational Fluid Dynamics (CFD) simulations and strategies for 3D-1D coupling between the embedding domain and the embedded structures. Finally, we outline some future perspectives aimed at enhancing accuracy, reducing computational time, and incorporating advanced modeling and boundary condition strategies to further refine the framework’s capabilities.

## 1. Introduction

In the rapidly evolving field of computational fluid dynamics (CFD) and simulation sciences, the precise and accurate representation of tubular structures, including those embedded within complex environments, emerges as a crucial element across a broad spectrum of applications. These applications span diverse fields, from detailed biophysical interactions within root-soil systems, as evidenced by previous studies [1, 2, 3, 4, 5, 6], to the exploration and biomimicry of ant nest architectures and their tunneling strategies [7, 8]. They extend further to the biomedical realm from the modelling of flows in blood vessels [9, 10, 11] and within lung airways [12, 13] to understanding tissue perfusion dynamics [14, 15, 16, 17, 18, 19, 20]. The relevance of (embedded) tubular structures also extends to industrial applications, for example the design and optimisation of heat exchangers [21], the efficiency of oil extraction processes at the scale of wells and reservoirs [22, 23], and safety considerations in tunnel ventilation systems [24] through to the engineering complexity involved in ensuring the stability and integrity of pipeline-soil systems [25, 26].

On the other hand, recent developments in imaging techniques (such as synchrotron or laboratory x-ray CT [27, 28, 29], neutron CT [30], and Multifluorescence High-Resolution Episcopic Microscopy [31]), coupled with novel segmentation and image processing methods [32, 33], have provided unprecedented access to realistic structures of tubular objects down to the finest length scales. Alternatively, when the required imaging resolution is lacking, it is possible to algorithmically generate biomimetic and morphologically-realistic synthetic tubular objects algorithmically [34, 35, 36]. This availability of structural data from imaging or synthetic sources presents both challenges and opportunities. Combining such data with mathematical models of physical processes provides the opportunity to interrogate the link between structural and functional relationships [37]. However, this relies on robust tools to extract the structural features from the raw imaging data [33].

Despite this opportunity, the pathway from obtaining three-dimensional imaging, synthetic data, or computer-aided design (CAD) models of (embedded) tubular objects to generating meshes that are ready for numerical simulations presents significant hurdles. This process is often labor-intensive and fraught with challenges, particularly when it requires the transition from raw data to refined, simulation-ready formats. Additionally, when these models are combined with finite element methods (FEM) to perform relevant simulations such as computational fluid dynamics (CFD) and advection-diffusion-reaction simulations, followed by the need for careful post-processing, the whole process becomes much more complex. This issue becomes even more challenging when trying to complete all these steps within a single open-source platform. In fact, the absence of seamless integration between various stages can make workflows more complex, particularly because of the format incompatibilities among different tools. Moreover, implementing an open-source plateform will increase the transparency and reproducibility of the research and will accelerate innovation within the scientific community.

Consequently, it’s crucial to develop a comprehensive highly-automated open-source pipeline that integrates the various steps of the process into a single framework. This unified approach would streamline workflows and promote the use of advanced simulation techniques across a wide range of applications. Such a workflow would mitigate the labor-intensive work associated with processing geometries and enabling the automated analysis of large datasets. This is particularly pivotal in fields such as the biomedical sector, where the capability to efficiently handle pre-clinical data (*e.g.* from animal models) and/or patient-specific data can significantly enhance diagnostic and therapeutic strategies on large multi-object datasets.

This paper is structured as follows: section 2 delineates our methods, starting with a comprehensive review of the software choices that underpin our pipeline and highlighting the interoperability of software packages and the automation strategies employed to streamline these processes. The different compartments are then described including the surface mesh processing from network-based and image-based inputs, the skeletonisation of tubular structures, the boundary condition automation, the mesh generation and conversion, the Finite element methods for the mathematical models integrated in the framework and finally the post-processing and visualisation of the simulation output. We showcase various examples in section 3 to demonstrate the applicability and efficacy of our framework across a range of scenarios, underscoring its potential to significantly advance the field of (embedded) tubular structure simulations. Finally, a list of potential enhancements is outlined in section 4, delineating the path for future advancements before concluding the paper in section 5.

## 2. Methods

### 2.1. Software/packages choices

In developing our pipeline, we have strategically selected a combination of powerful general-purpose opensource software packages to ensure efficiency and flexibility across various stages of our workflow. It is worth mentioning that all the software and packages were invoked through a unified *MATLAB* script to maximise automation capabilities.

The first part of the pipeline generates and processes surface meshes from different inputs (3D imaging, CAD models, synthetic/skeletonised Networks), using an open-source *MATLAB* [38]-based toolbox called *GIBBON* (The Geometry and Image-Based Bioengineering add-On)[39]. *GIBBON* includes a wide range of CAD, geometry and image processing tools. This toolbox also enables interfacing with free open-source softwares such as *TetGen* [40] and *Geogram* [41] that both possess high-quality meshing capabilities.

The second part of our pipeline involves extracting the centre-lines of the tubular structures. We integrate two versatile skeletonisation algorithms, *VesselVio* [42] and *TreeSkel* [43], that have been used respectively on vascular and airways datasets.

For the automatic labeling of surface and volumetric mesh generation and conversion in the third and fourth segments of the pipeline, we rely on the features offered by *GIBBON* and *meshio* [44, 45]. The *meshio* Python library is designed for handling mesh data. It provides functionalities for reading, writing, and converting mesh files in various formats commonly used in computational simulations and visualisation. There are numerous open-source finite element software options which have been developed to solve free fluid flow and Darcy-scale porous-medium flow problems as well as coupled partial differential equation (PDE)-based transport models, such as *Dumu^x^* [46, 47] and *FEBio* [48, 49] (a detailed list can be found in [50]). We opt for *FEniCS* [51] [52] [53], which is a general-purpose open-source software framework designed to streamline the process of discretising PDE using the finite element method. *FEniCS* leverages the capabilities of the Unified Form Language (UFL) [54] and the *FEniCS* Form Compiler (FFC) [55] to automatically produce optimised low-level C++ code for evaluating equations expressed in finite element variational forms. Additionally, it provides interfaces to PETSc and Trilinos for linear algebra computations. To use *FEniCS*, the user is required to provide the high-level variational form of the differential equation and doesn’t have to perform any coding for the discretised form at the level of the cell and element scale. *FEniCS* is also well-suited for advanced solid mechanics simulations, particularly when integrated with *MFront* [56, 57, 58], allowing therefore a coupling between flow and mechanics.

Finally, for post-processing purposes, ParaView [59, 60] (built on the Visualisation Toolkit (VTK) library [61]) stands out as a comprehensive visualisation tool, facilitating intuitive exploration and analysis of simulation outcomes. Its broad array of features and compatibility with multiple data formats render it an essential resource for extracting valuable information from simulation results.

Figure 1 illustrates the interoperability of the different softwares/packages used in the Tube2FEM pipeline. Next, we describe the components of this workflow in turn.

**Figure 1:**
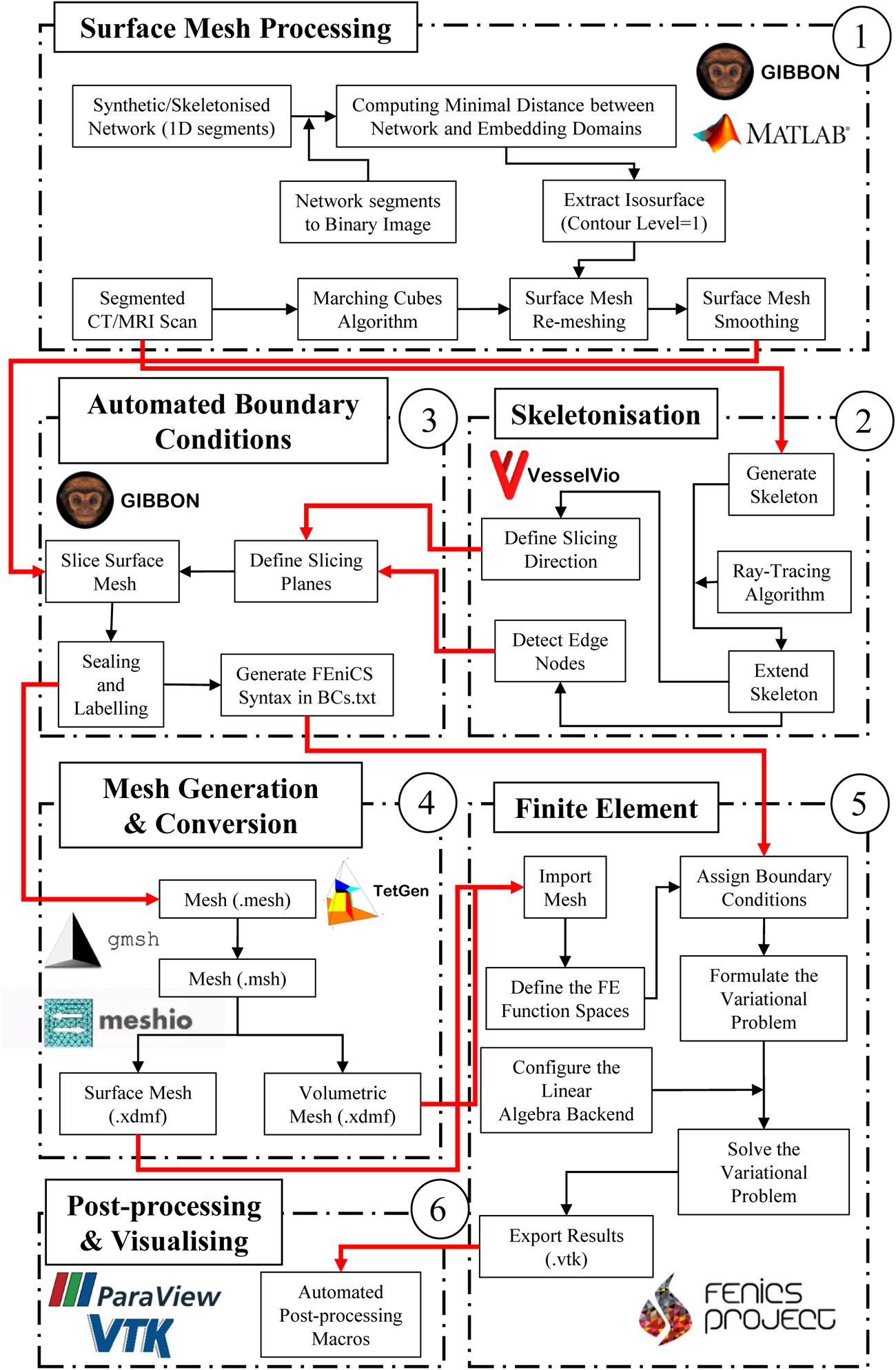
Interoperability of the software/packages used in the Tube2FEM pipeline.

**Table 1:**
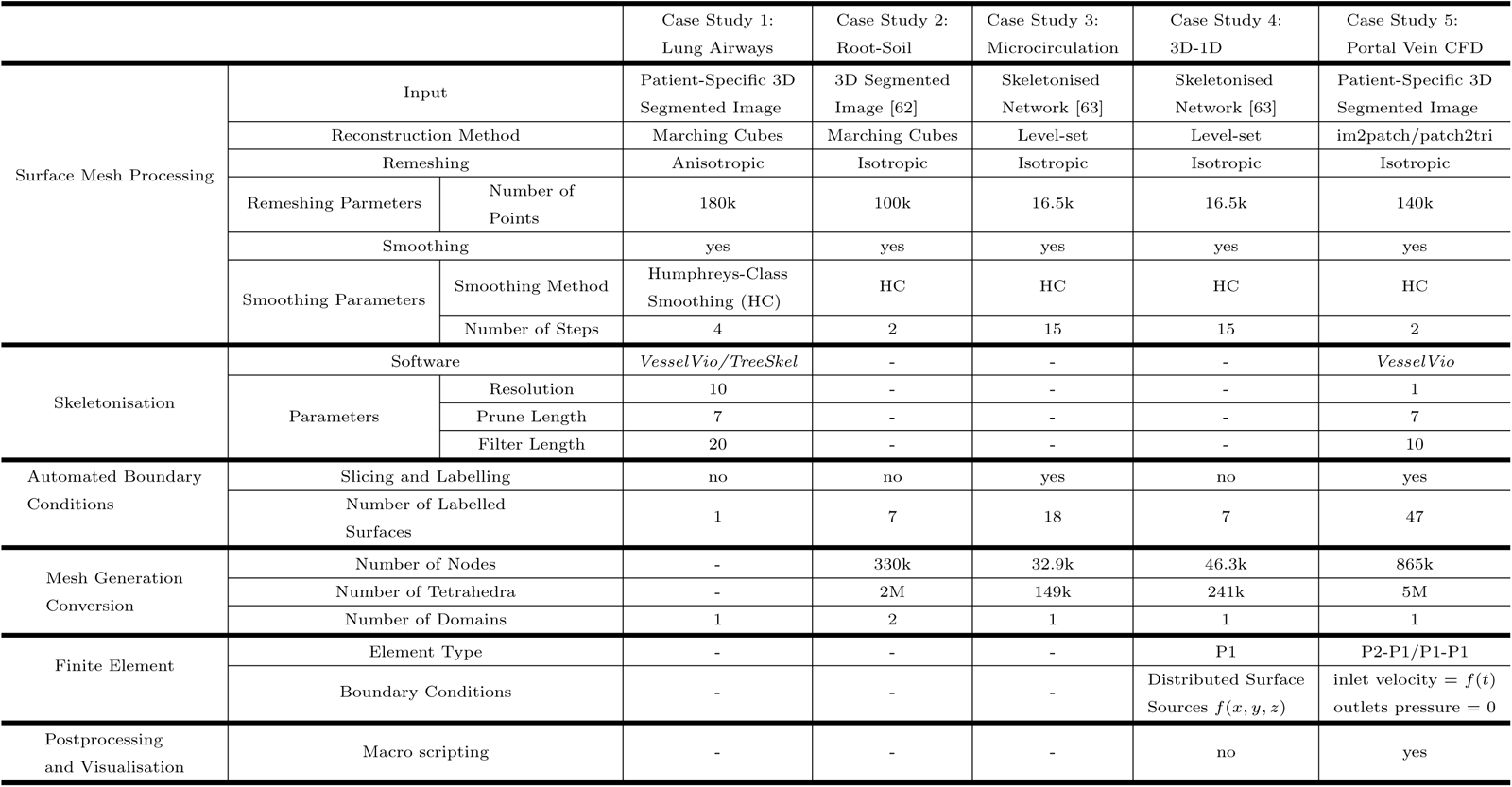
Summary of key parameters and method specifications in Tube2FEM for the five case studies presented in this work.

### 2.2. Generating a watertight surface mesh

This workflow is intentionally flexible and can handle different inputs, including segmented 3D images issued from different imaging tools (X-ray Computed tomography (XTC), Optical Coherence Tomography (OCT), Magnetic Resonance Imaging (MRI), *etc*) and synthetic/skeletonised graph networks.

For the first type of input, namely, the segmented 3D image, the creation of the surface mesh can be efficiently accomplished by employing the computer graphics marching cubes algorithm [64]. We use the implementation of the Marching Cube from the *MATLAB* file exchange [65] for this purpose. It is also possible to use the *GIBBON* functions im2patch and patch2tri to get a watertight surface mesh.

For the alternative input form, namely skeletonised or synthetic graph networks, continuous surface models are obtained by using level-set images. The procedure for creating level-sets typically includes 3 steps. The initial step involves embedding the spatial data within an image domain. The second step entails computing a distance function between the spatial data nodes and the image’s voxel grid. The last step involves using the distance function to produce a (signed) level-set image. From there, continuous surface geometries can be obtained through the calculation of isosurfaces. A detailed description of this method can be found in [10] and an implementation of it already exists in *GIBBON*. Based on the graph network’s size and the desired level of precision, calculating the distance function between the network nodes and the image voxel grid can become significantly time-consuming. To address this, we enhance the efficiency of the function by creating a binary image from the graph network, enabling the distance map calculation to occur only between a dilated version of this binary image and the nodes of the graph network.

### 2.3. Surface Mesh processing

The watertight surface mesh, outlined in the previous section, exhibits voxel artefacts as a result of the used voxel-based methods. A re-meshing step is therefore required to eliminate these artefacts. We use *Geogram* which is an external library in *GIBBON* to remesh the triangulation defined by the mesh faces (F) and the vertices (V) [41]. In particular, the code *Vorpalite* is used and optimises the Voronoi diagram from the point of view of sampling regularity. The resulting mesh consists of a near-isotropic distribution of triangles that transforms the stair-shaped surface patterns to a smoother appearance. We follow that by using the patchSmooth function available in *GIBBON* to complete the smoothing operation. In particular, we use the option of “Humphrey’s Class” that preserves best the overall volume of the watertight surface mesh.

### 2.4. Skeletonisation

In order to run physics-based simulations, it is necessary to assign boundary conditions on the edges of the network (for example to impose fluid pressure, velocity or solute concentration conditions). This requires creating flat surfaces at the terminal branches of the network and labelling them. While this task can be done manually for small networks, it becomes cumbersome and time consuming for medium and large networks. Thus, automating this process is crucial to make this pipeline viable for a broader spectrum of applications. One approach to automation involves baselining the cutting and labelling process around the terminal branches of a network’s centre-line volume. Here we describe the step in the pipeline which creates the centre-line for the tubular volumes.

We assume that the centre-line of the vessels is equivalent to the skeleton obtained by skeletonisation algorithms in the context of image-processing. An easily accessible open-source tool for 3D tubular-dataset analysis is therefore needed. Moreover, to facilitate automation, this tool must be designed to operate as a standalone executable application, negating the need for graphical user interface (GUI) interaction. VesselVio [42] meets these requirements and is compatible with both Windows and MacOS. It also provides high-level Python libraries, just-in-time compilation, and parallel processing for rapid and detailed feature extraction techniques. This latest feature allows the transformation of the skeleton image to a graph format. For Linux users, the *TreeSkel* package [43], employing the minimum cost paths algorithm [66], offers an alternative solution for skeletonisation.

The smoothed surface mesh was converted back into a binary image, due to the potential alterations introduced by the remeshing and smoothing operations on the original binary image. *VesselVio*, or alternatively *TreeSkel*, was then used to derive the skeleton of the network. Furthermore, plugins were created to import the graph format output into MATLAB. However, when *VesselVio* is used, overlaying the skeleton on the surface mesh reveals a noticeable “retraction” effect, where the skeleton’s edge points do not extend fully to the surface mesh. This is expected since *VesselVio* employs a custom implementation of a widely used medial axis parallel thinning algorithm [67]. This retraction poses a challenge as the edge points are crucial for guiding the slicing and labeling of surfaces to create boundary conditions. To address this issue, a Ray-Tracing algorithm is employed to elongate the edge sub-segments until they intersect with the surface mesh. These intersection points are then integrated into the skeleton, serving as the new edge points.

### 2.5. Anisotropic mesh to enhance computational accuracy and efficiency

Prior to employing the edge points and terminal branches of the skeleton to guide the cutting and labeling of the surface mesh, a crucial step in the pipeline is necessary, particularly for networks composed of vessels with a wide variation in diameters. Users may encounter a challenge in selecting an appropriate element size for the isotropic mesh. Choosing a small element size better preserves the morphology of the smaller tubular structures, yet results in excessive meshing of the larger ones, leading to a significant increase in the computational time required for the physics-based simulations. On the other hand, opting for a larger element size can lead to inaccuracies in representing the morphology of smaller tubes, along with a deficiency in the number of elements on the edge surfaces where boundary conditions are applied. To address this issue, employing an anisotropic meshing approach, where the degree of anisotropy is guided by the tubes’ radii, is required. A minimum distance is thus calculated between the nodes of the surface mesh and the skeleton nodes using the minDist function found in *GIBBON*. This data is then fed to *TetGen* [40], an external library interfaced by *GIBBON*, which generates an anisotropic surface mesh. In this resulting mesh, smaller vessels are finely detailed, whereas larger vessels are meshed more coarsely.

### 2.6. Slicing and labelling to enable boundary condition assignment

To assign boundary conditions, it is essential to generate terminal cross section surfaces through the tubular structure. For broader applicability, these surfaces should be flat, particularly to provide sufficient flexibility for use of vectorial entities, and they must contain a sufficient number of mesh elements to avoid numerical instabilities. This can be accomplished by slicing the surface mesh’s terminal branches along their edges and subsequently re-meshing to ensure the entire mesh remains watertight. Thus, accurate slicing operations are required at each terminal branch.

A slicing plane is defined by a point and a normal vector. For each slicing operation performed on the tubular geometry, the point of origin is an edge point of the skeleton, and the normal vector corresponds to terminal sub-segment of the skeleton. However, since a slicing plane is infinite, it might indiscriminately affect other sections of the mesh. To enhance the accuracy of this operation, a Boolean constraint is introduced, ensuring that only interconnected elements nearest to the skeleton edge point are impacted. This is achieved by using the minDist function to apply the Boolean condition, followed by the triSurfSlice function for executing precise slicing on the targeted branch. Both functions are part of the *GIBBON* toolbox.

Following that, the now open surface mesh is re-sealed by re-meshing the terminal branches using a series of functions, the most important of which is regionTriMesh3D (available in *GIBBON*). Each newly formed surface is assigned a distinct label, and an automatically generated text file, containing essential details about these surfaces, is created. This text file can be directly communicated to *FEniCS* to apply the boundary conditions.

### 2.7. Volumetric meshing

The labelled surface mesh is now prepared for a second processing by *TetGen* [40] to create a volumetric mesh, ready for physics-based simulations. We input specific parameters into *TetGen*, including meshing options, faces, nodes, face labels, and the number of regions. The runTetGen command in *GIBBON* is then used to produce a tetrahedral mesh. The output includes the tetrahedral elements, element IDs, faces, face IDs, and the coordinates of the nodes. It is also possible to convert the tetrahedral elements to hexahedral ones using the tet2hex function.

### 2.8. Mesh exporting and conversion

Different finite element (FE) software packages require various mesh formats. The FE software used in our pipeline is *FEniCS*, which supports a limited number of formats, including .xml and .xdmf. To accommodate this, we have developed a function capable of converting the output from a *TetGen* mesh into a version 2 ASCII *Gmsh* file (.msh format). Subsequently, a Python script employing the *meshio* library is used to convert the .msh file into an .xdmf format, ensuring compatibility with *FEniCS*.

### 2.9. Mathematical Models

We integrate physics-based mathematical models into this workflow, which are commonly utilised in simulating processes that involve (embedded) tubular objects. The first model is the incompressible Navier-Stokes model to simulate the fluid dynamics in the tubular objects. The second model is to simulate the advection-diffusion-reaction of specie(s) (solutes, oxygen, drugs, nanoparticles, *etc*) in the embedded or embedding domain. While these are specific examples, the broad framework can be easily adapted to other physical models (e.g. solid mechanics).

#### 2.9.1. Computational fluid dynamics

We assume that the fluids of interest are non-compressible Newtonian fluids and so fluid flow can be described via the incompressible Navier-Stokes equations, written in the form:

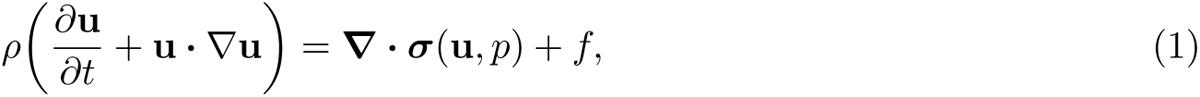

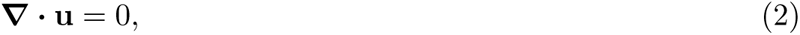

where *f* is a given force per unit of volume, **u** is the velocity of the fluid, *p* its pressure, *ρ* its density and ***σ***(*u, p*) is the stress tensor. For Newtonian fluids, the stress tensor is given by

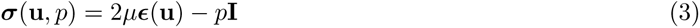

where **I** is the identity matrix and ***ɛ***(**u**) is the strain-rate tensor:

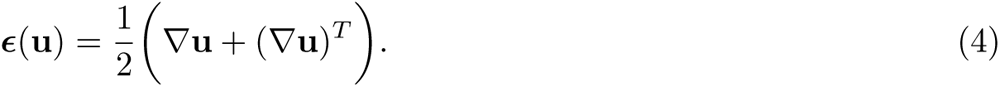

The Navier–Stokes equations exhibit non-linearity, transience, and a non-trivial coupling between pressure and velocity. While coupled methods exist to solve the system of equations, it is more popular to decompose the problem into several more manageable equations. The initial splitting scheme was proposed by Chorin [68] and Temam [69]. This method was further refined by Goda [70], who introduced a velocity correction step, and his approach became widely recognised in the literature as the Incremental Pressure Correction Scheme (IPCS). Despite the existence of alternative splitting schemes, a thorough comparison presented in [71] indicates that, in terms of both efficiency and accuracy, the Incremental Pressure Correction Scheme (IPCS) generally outperforms other methods. Moreover, the IPCS method is relatively more straightforward to implement when contrasted with alternative splitting schemes. It consists of solving three equations:

- the tentative velocity step (a convection–diffusion–reaction equation),
- the pressure-correction step (a Poisson equation),
- the velocity update step (a projection).

We customise the implementation of this method provided in the *FEniCS* tutorial [72].

#### 2.9.2. Species transport in the embedded or embedding domains

We include general-purpose transport of chemical species, described using an advection-diffusion-reaction equation,

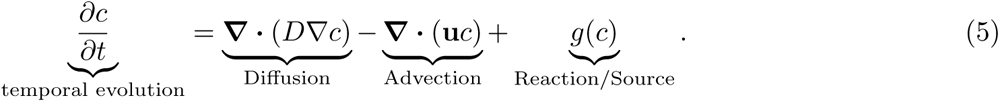

Here the temporal evolution of a chemical species of concentration *c* is linked to its spatial evolution through diffusion, advection by a vector field **u** and/or a reaction within the domain. It is worth noting that this equation is usually coupled (weakly) with the aformentioned Navier-Stokes through the velocity field **u** especially when the advection effects are important.

#### 2.9.3. Mixed-dimension models: 3D-1D coupling

Our framework is generic and suitable for flow systems with high, intermediate, or low Reynolds numbers. However, there are numerous situations where it is practical to consider steady-state flows at low Reynolds numbers, such as blood flow in networks of small blood vessels and plant root systems. In these cases, it is reasonable to disregard changes in flow velocity and inertial effects. This is reflected mathematically by eliminating both the time-dependent and inertial terms in the Navier-Stokes equations, leading to the use of the simpler Stokes flow equations. Moreover, in such situations, a one-dimensional model of the tubular structure is often adequate to capture the essential aspects of flow physics and is often used in the literature [14, 17, 15]. Nevertheless, when the focus is on how the 1D tubular objects interact with their 3D surrounding environment, a significant challenge emerges due to the mixed-dimensional (3D-1D) nature of the problem. In fact, various numerical methods vary in their approach to bridging the dimensional incompatibility between network and embedding domain by distributing source terms. The contribution of the source term within the encompassing bulk medium can be represented through line source terms [73, 74], surface source terms [75, 76], or volume source terms [18, 4]. In many of these methods, it is assumed that the diameters of the network branches are significantly smaller than the dimensions of the surrounding domain. This facilitates a simpler mesh creation process that results in an overestimation of the bulk volume by covering the space occupied by the tubular structures. Additionally, due to the elimination of their physical structure, the tubes no longer offer a resistance to flow in the embedding domain. Finally, as these mixed-dimension models typically represent the network by a series of finite cylinders, inaccuracies arise at junctions where source terms should be distributed.

Most of these problems in 3D-1D coupling settings can be avoided if the exact network surface mesh is known. Our framework allows for the building of such continuous surface meshes from networks of 1D segments, as shown in section 2.2. Consequently, simulations conducted on the network of 1D segments can be projected onto the network surface mesh using an minimal distance-based approach between the nodes of the 1D segments and those of the surface mesh. Then, using the *TetGen* [40] interface with *GIBBON*, a conforming mesh that integrates the embedded system and its surrounding environment can be created. The interface conformity between the two domains allows the direct use of the distributed source terms on the network surface without the necessity of extrapolation.

### 2.10. Scientific Visualisation and Post-processing

Geometries and meshes are visualised in *MATLAB* using the *GIBBON* open-source toolbox, whereas the simulation results are exported in *.vtk* format and are visualised and post-processed using *ParaView* [59, 60] (built on the Visualisation Toolkit (VTK) library [61]). For automation purposes, the use of the *ParaView* Graphical User Interface (GUI) must be avoided. For that, generalised macros were written and called from *MATLAB* through a Command-Line Interface (CLI).

## 3. Results and Discussion

This section highlights five case studies that comprehensively illustrate the capabilities of the Tube2FEM framework. Particularly, we present instances of tubular geometries corresponding to three distinct applications (lung airways, root-soil systems, micro/macro circulation), each emphasising various aspects of the framework, including surface mesh reconstruction, meshing, CFD simulations, 3D-1D approaches, visualisation, *etc*. We note that all the case studies presented in the upcoming section are automated, with each having all of its commands centralised within a single *MATLAB* script. Finally, Table 1 summarizes all the key user-defined parameters, method specifications and some of the output results for all of the 5 case studies. With the exception of the cases featuring patient-specific data, all the codes are made available on GitHub at: https://github.com/CheikhSleiman/Tube2FEM

### 3.1. Case study 1: Radius-dependent anisotropic mesh generation of lung airways

For our initial example, we employ Tube2FEM to generate simulation-ready meshes from segmented lung airways acquired using medical CT scans. The primary difficulty with this type of tree-like tubular geometry is its topological complexity, since it contains more than 10 branching generations, and there is significant variation in diameter between the smallest and largest airways. Indeed, the challenge stems from choosing a mesh element size that maintains the airways’ morphology without leading to increased computational time for the simulations. Tube2FEM addresses this issue by employing a radius-dependent anisotropic meshing approach. The process begins with the segmented lung airways volume, as depicted in Figure 2(a), from which the tubular structure’s centre-line is determined using skeletonisation algorithms, as detailed in section 2.4. Additionally, the surface mesh is generated from the 3D image using the marching cubes algorithm or similar techniques, as described in section 2.2. However, this mesh initially exhibits voxel artifacts, necessitating further processing steps such as remeshing and smoothing, discussed in section 2.3. Subsequently, the processed surface mesh is overlaid with the skeleton, as shown in Figure 2(c). The next phase involves projecting the centre-line radii field onto the surface mesh, based on the minimal distance criterion outlined in section 2.5. This projection creates a radius-dependent weighting map that guides the mesh’s anisotropy, shown in Figure 2(d). The final comparison between the preprocessed surface mesh, the isotropically remeshed and smoothed version, and the anisotropic mesh is presented in Figure 2(e), (f), and (g) respectively.

**Figure 2:**
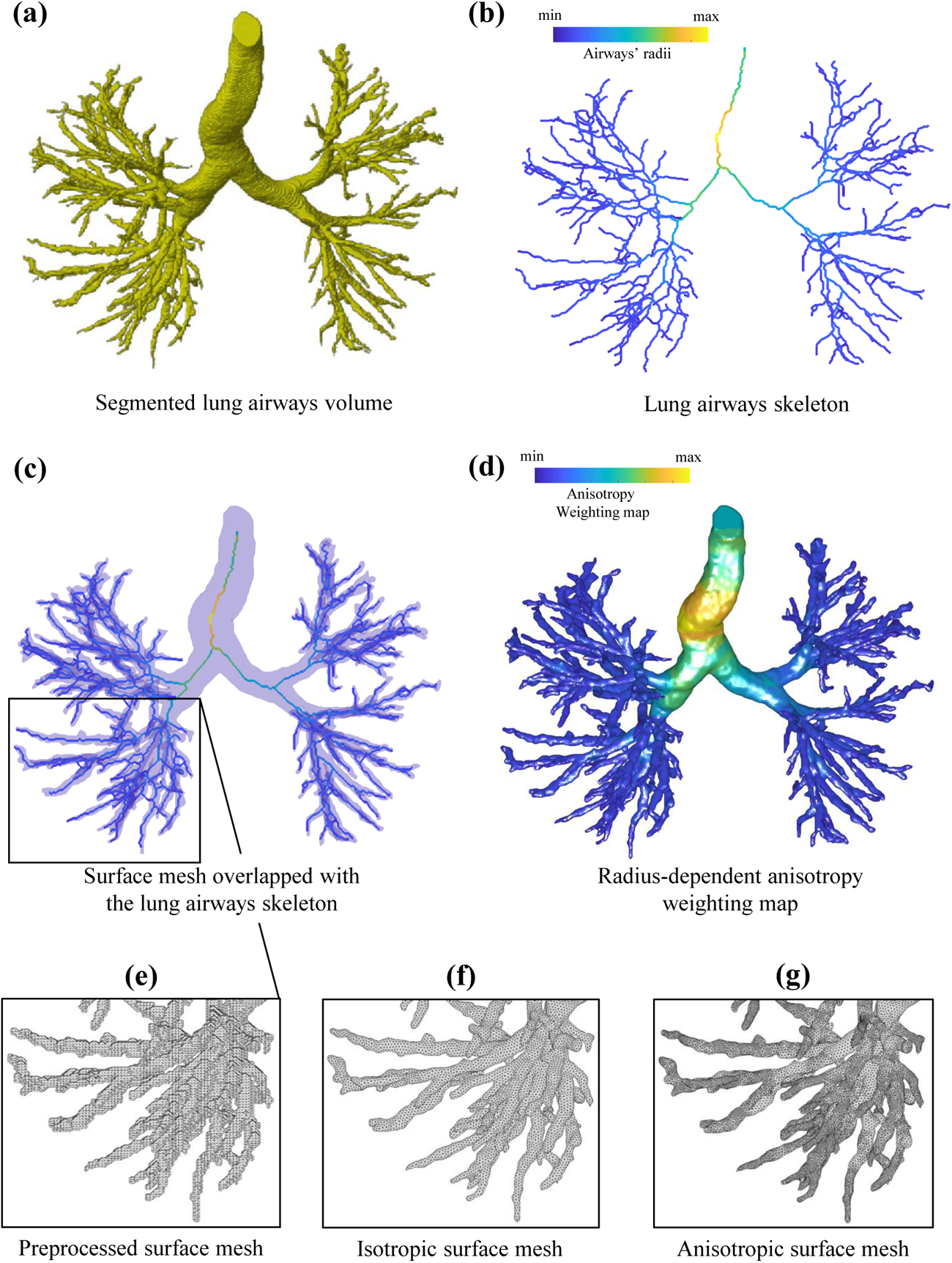
The Tube2FEM procedure for creating radius-informed anisotropic meshes for a lung airway tubular geometry acquired using CT imaging.

### 3.2. Case study 2: Multi-domain mesh generation of a root-soil system

In this case study, Tube2FEM is used to generate a multi-domain, simulation-ready mesh of a root network embedded within a soil medium. The root network data, previously examined in [4], is accessible online in Gmsh (.msh format) as referenced in [62]. To optimise the effectiveness of this procedure, we perform several operations to convert this format into 3D imaging (.tiff format), which is commonly adopted in the literature. We then use this transformed dataset illustrated in Figure 3(a) as the input for our procedure. As in the previous case study, we extract the surface mesh using the marching cubes algorithm or equivalent techniques, as outlined in section 2.2. Remeshing and smoothing processes are then carried out to eliminate voxel artifacts and achieve a desired mesh density, as illustrated in Figure 3(b) and (c). Indeed, in order to produce reliable results in hydraulic root-soil simulations, the mesh needs to be carefully locally refined at the interface. This is particularly important as drying soil around the roots creates large and highly localised pressure gradients. The effective way to capture these gradients is through local mesh refinement at the interface [4]. A soil medium is then built around the root mesh using *GIBBON*’s triBox function, and both media are inputted into *TetGen*, as detailed in section 2.7, to create a multi-domain tetrahedral mesh. Figure 3(d) shows both domains, featuring a mesh density gradient from the interface to the boundaries of the soil domain.

**Figure 3:**
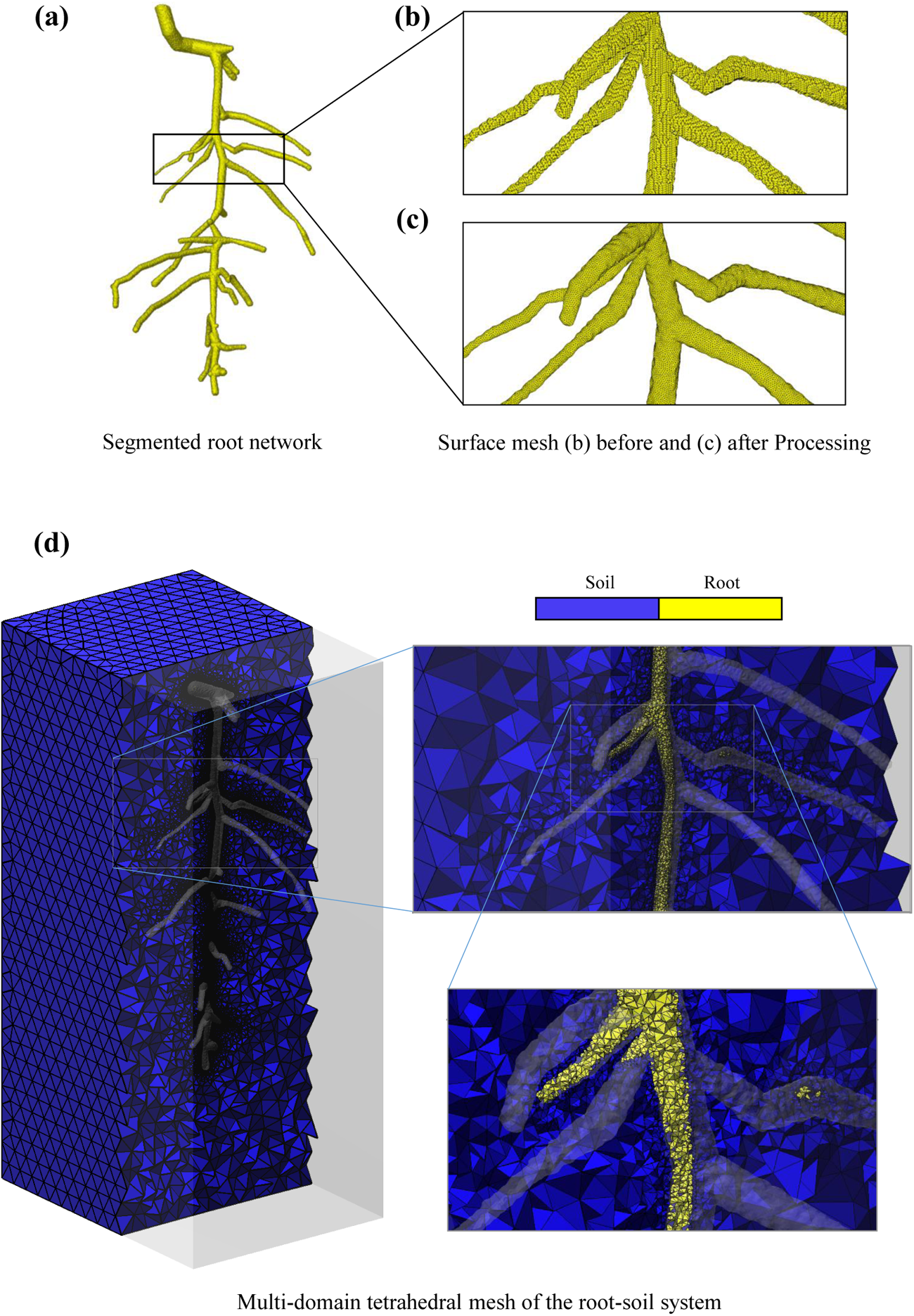
The Tube2FEM procedure for creating a multi-domain tetrahedral mesh of a root-soil system, based on MRI imaging

### 3.3. Case study 3: Generating surface and volumetric meshes of a microcirculation blood Network

This case study demonstrates the ability of Tube2FEM to generate simulation-ready labelled surface and volumetric meshes from networks of 1D tubular objects. These networks can either be synthetic (for example, via generative algorithms) or result from skeletonisation processes applied to imaging datasets. While the procedure is general, we apply it to a widley-adopted microvascular network dataset used in the literature to explore microcirculatory blood flow and transport processes, studied initially in [63].

The level-set method, detailed in section 2.2, is used here to create a watertight surface mesh. The starting point is the vascular network centre-line over which the vessel radii field is presented, as illustrated in Figure 4(a). The second step entails embedding the spatial graph within an image domain and then computing a distance function between the centre-line nodes and the image’s voxel grid. This method can become significantly time consuming if the image domain is not carefully chosen. To address this, we create a binary image from the graph network as illustrated in Figure 4(b). Several image dilation operations are then executed to establish the required image domain. The signed level-set image, shown is Figure 4(c) can be interpreted as the normalised minimal distance map between the centre-line nodes and the dilated binary image. The isosurface, corresponding to level-set intensity = 1, can be extracted from the level-set image. The resulting surface mesh, which has been remeshed and smoothed to eliminate artifacts, is displayed in Figure 4(d). The following step involves applying the slicing and labeling algorithm, described in section 2.6, to assign boundary conditions. This is achieved by overlaying the centre-line network with the processed surface mesh and identifying the edge nodes and the directionality of the edge segments. Finally, having the watertight labelled surface mesh illustrated in Figure 4(e), it is possible now to generate the tetrahedral mesh using the *TetGen* interface in *GIBBON*. The volumetric mesh in shown in Figure 4(f).

**Figure 4:**
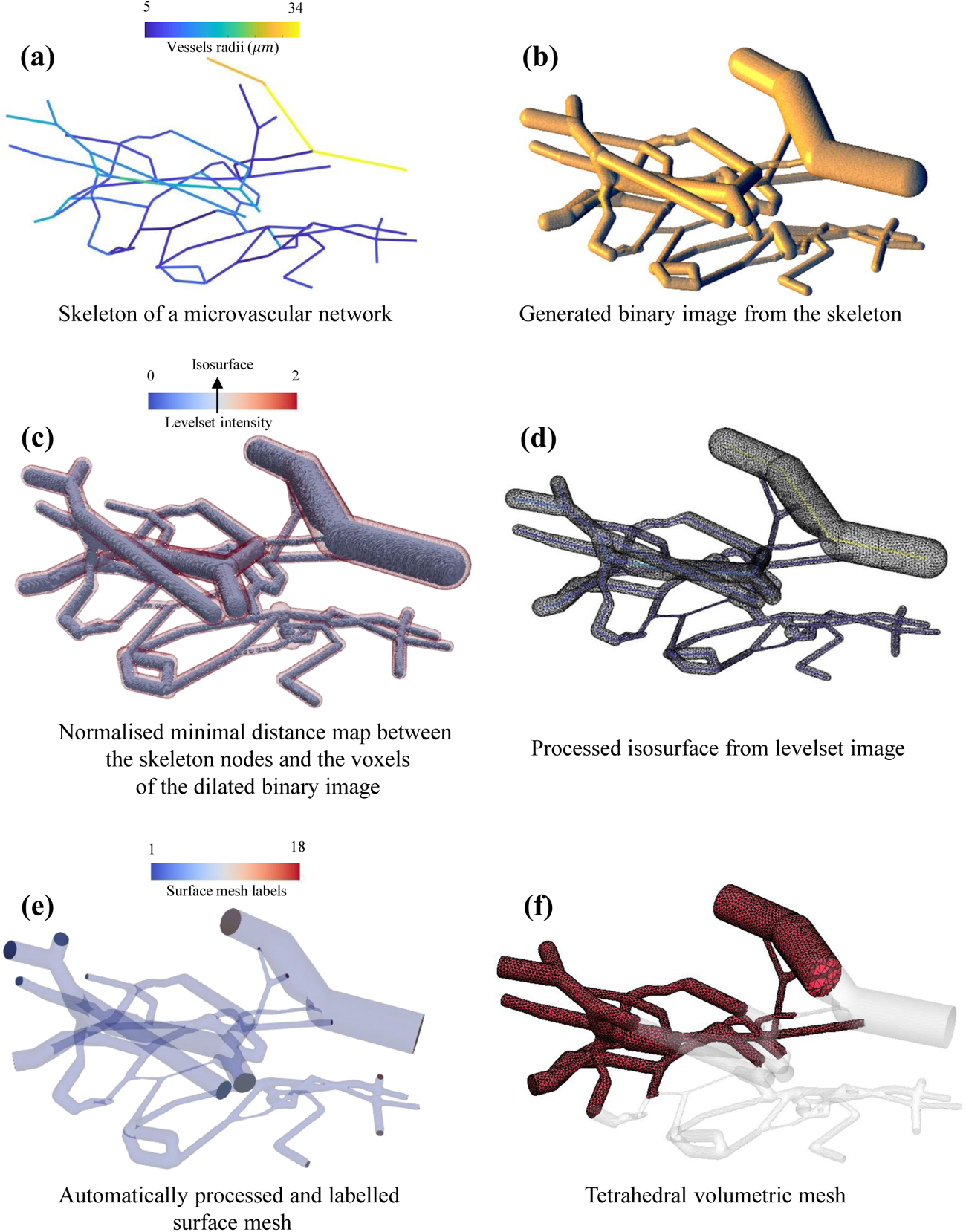
The Tube2FEM procedure for generating labelled surface and volumetric meshes from a skeletonised microcirculation tubular geometry [63].

### 3.4. Case study 4: 3D-1D coupling strategy to simulate tissue perfusion

This case study aims to demonstrate the ease of 3D-1D coupling between an embedding tubular network and its surrounding domain using Tube2FEM. For convenience, we apply this procedure on the same graph network used in case study 3 and the same reconstructed surface mesh which was created using the levelset method. The final goal is to simulate the extra-vascular solute transport driven by a heterogeneous distribution of solutes within the vascular network.

In this specific scenario, a 1D representation of each vessel suffices to capture the flow physics. This renders the computational time cost of the simulated intravascular processes very cheap in comparison to a 3D case scenario. Consequently, the initial step in this process involves simulating solute transport across a network of 1D vessels. This simulation can be conducted using methods such as Finite Element, Finite Volume among others [14, 15]. However, when attention shifts to the interactions between the 1D tubular objects and their 3D surroundings, a considerable challenge arises from the mixed-dimensional (3D-1D) nature of the boundary. As discussed in section 2.9, different numerical methods adopt various strategies to overcome the dimensional mismatch between the network and the embedding domain by distributing source terms (line, surface or volume sources terms). For this case study, we distribute the source terms across the surface, since Tube2FEM is capable of reconstructing the exact surface mesh of the vascular network, as demonstrated in Case Study 3. To achieve this, we proceed to overlap the simulation results on the centre-line with the reconstructed surface mesh as illustrated in Figure 5(b). Then the solute field on the network of 1D vessels is projected on the mesh using a minimal distance computation between the centre-line nodes and the mesh ones. This results in the surface field shown in Figure 5(c). Following this, we create the surrounding tissue volumetric mesh using *TetGen*-*GIBBON* interface. In the input options for *TetGen*, we ensure that the volumetric tissue mesh should be hollowed out from the vascular domain, which is represented by the reconstructed surface mesh. We also ensure that the mesh density of the vascular surface mesh is sufficiently fine to more effectively resolve the volumetric mesh density gradient near the vascular-tissue interface. The result of these operations can be visualised in Figure 5(d). Additionally, we note that the internal surface of the volumetric mesh, created by the removal of the vascular domain, conforms to the reconstructed vascular surface mesh. Such conformity is essential for facilitating consistent interactions between the different domains. The volumetric mesh produced is first exported in the (.msh) format and subsequently converted to the (.xdmf) format using the *meshio* package. This latter format ensures compatibility with the *FEniCS* software, which is employed to carry out the Finite Element Analysis. We also export the solute field evaluated on the vascular surface mesh in a tabular format (*x_N_*, *y_N_*,*z_N_*, *c_s_*), where *x_N_*, *y_N_* and *z_N_*are the vascular surface mesh nodes coordinates and *c_s_* is the concentration of the solute on the node *N* . This table is then communicated to *FEniCS* and is used in assigning boundary conditions. Finally, we use the Poisson equation to model a simple steady-state solute diffusion process in the tissue as illustrated in Figure 5(e). We note that this procedure can be used for much more complex problems, and can be further developed, for instance to account for time-dependent intravascular flow variations, and two-way coupling.

**Figure 5:**
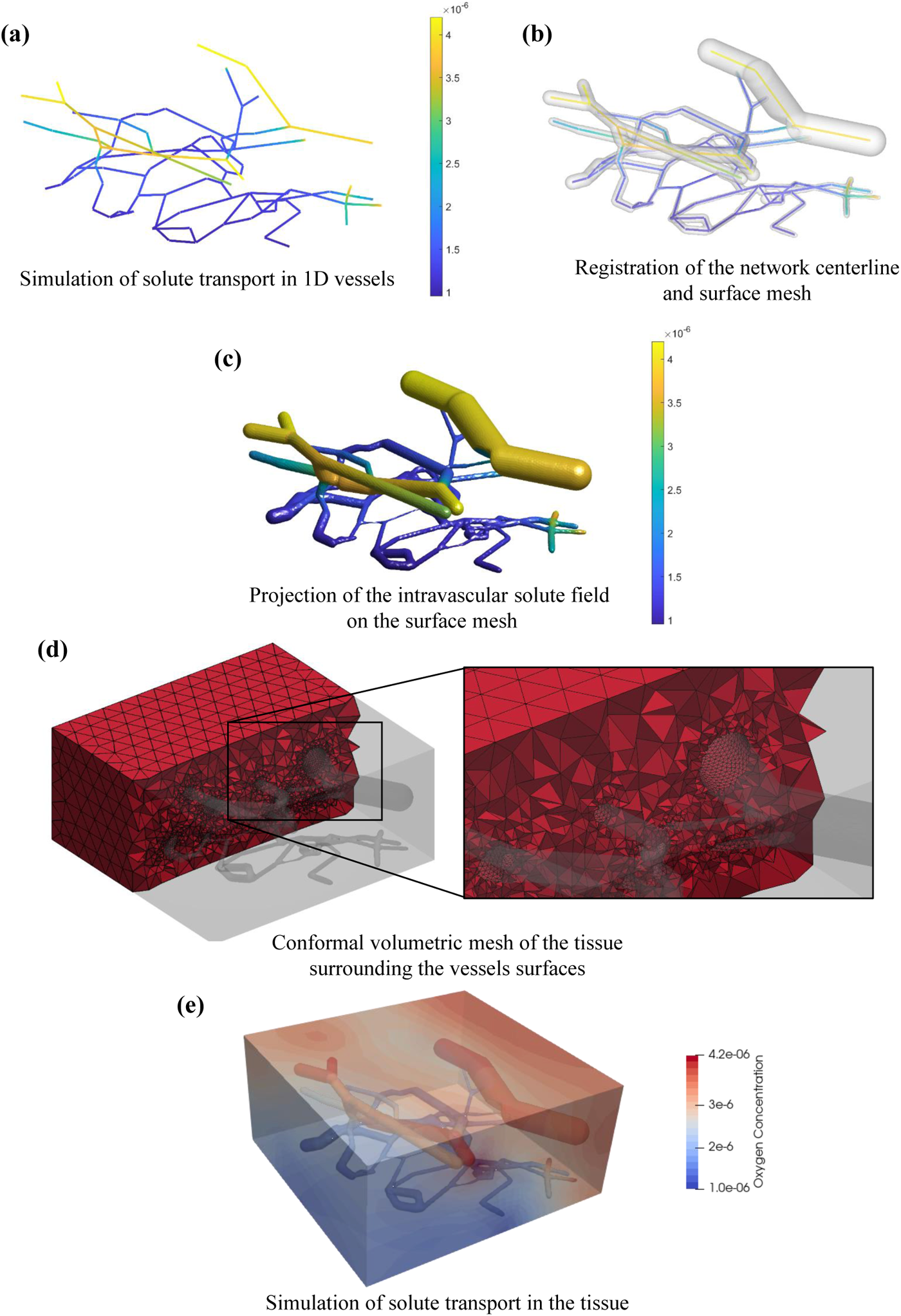
A 3D-1D coupling strategy enabling solute transport from a vascular network composed of 1D segments into the surrounding 3D biological tissue.

### 3.5. Case study 5: Detailed CT-to-simulation procedure for tubular geometries: application to portal vein network

This final case study is designed to showcase a comprehensive framework for conducting physics-based simulations on tubular geometries obtained via imaging. While the framework is broadly applicable, we demonstrate its use through a portal vein network derived from a patient’s CT scan. This segmented network is illustrated in Figure 6. Similarly to previous case studies, the portal vein surface mesh can be extracted using marching cubes algorithm or an equivalent method detailed in section2.2. The surface is then remeshed to a desired density and smoothed to get rid of geometrical artifacts. Following that, we call *VesselVio* software in *MATLAB* and skeletonise the binary image. The overlap of the resulting centre-line and the surface mesh can be seen in Figure 6(b). Subsequently, the skeleton edge nodes are identified, and when combined with the direction of the skeleton terminal sub-segments, they determine the slicing planes. The surface mesh after the slicing operations is visualised in Figure 6(c). The open geometry is then sealed by remeshing and labelling the open surfaces near the skeleton terminal branches. The detailed slicing, sealing and labelling procedures are detailed in section 2.6. In total, Figure 6(d) shows 47 labelled surfaces, 46 of which are created by sealing the sliced geometry and 1 being the vessels wall. Those labelled surfaces are crucial to assign boundary conditions for the physics-based models. For the Navier-Stokes (NS) simulation, we apply a time-dependent velocity to the inlet, zero pressure to the outlets and zero velocity to the vessel walls (no-slip boundary condition). Figure 6(e) illustrate the velocity streamlines at a specific time point, resulting from solving the NS problem. We then save all the the velocity fields at the different time steps and use them to inform the advection of the chemical specie in the advection-diffusion-reaction problem. Figure 6(f) shows the resulting concentration of the chemical field within the portal vein network. It is important the highlight that those simulation results are demonstrative to the capacity of the pipeline to solve complex flow and flow-related problem. The set of parameters used in those simulation doesn’t necessarily represent patient-specific physiological ranges.

**Figure 6:**
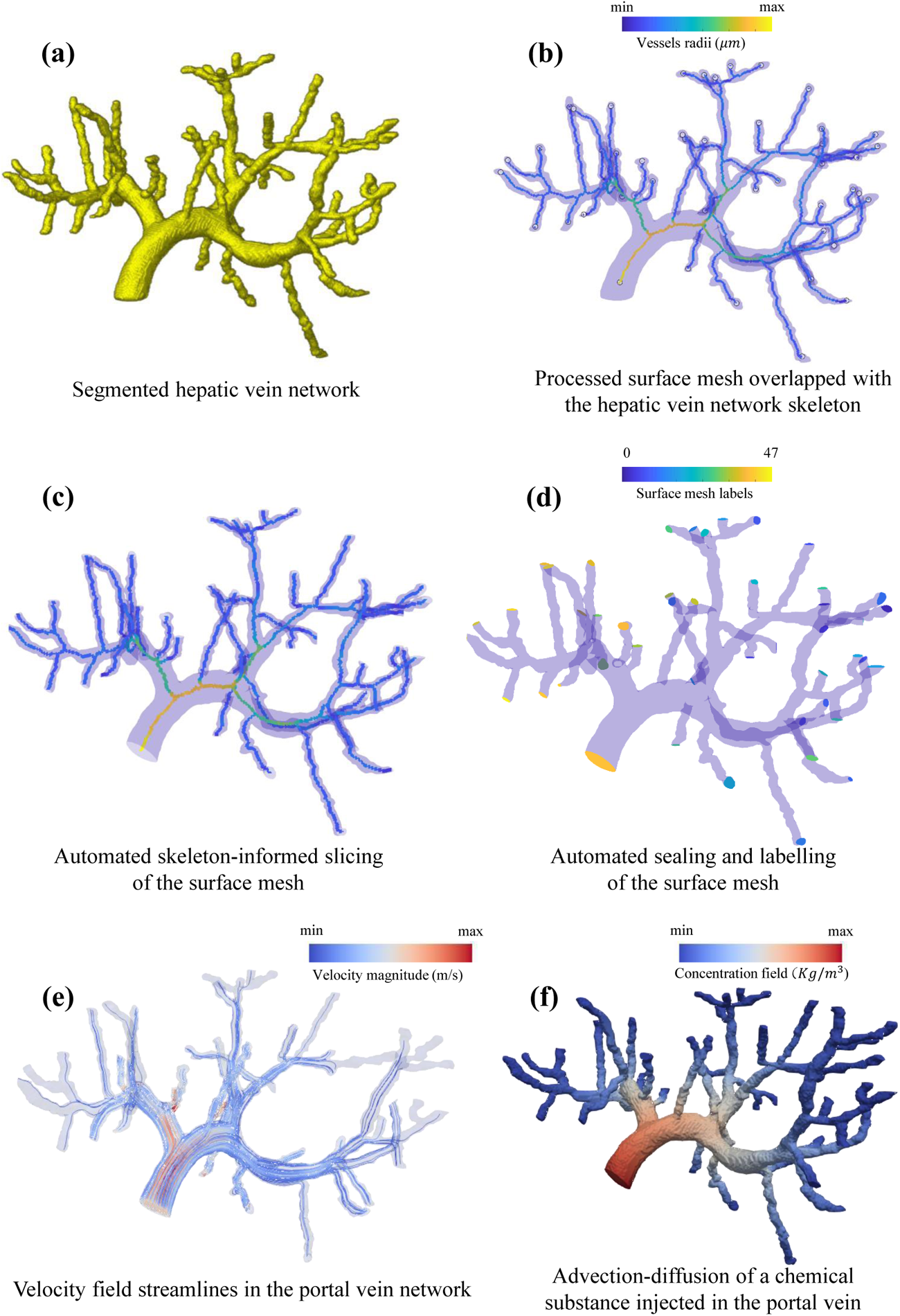
Image-based procedure detailing all the necessary steps to conduct CFD and advection-diffusion-reaction simulations from a segmented vascular geometry.

## 4. Perspective

Anticipating future advancements of our pipeline, we highlight critical improvements targeted at enhancing the accuracy and reducing the computational time required for simulating (embedded) tubular structures. The outlined perspectives below may be explored in future work to enhance the framework’s efficiency and technical capabilities.

- **High-Performance Finite Element Navier-Stokes Solver** *Oasis* is a Python-based high-performance finite element Navier-Stokes solver [77]. It utilises foundational components from the *FEniCS* project and is designed for addressing large-scale applications involving intricate geometries, particularly on massively parallel computing clusters.
- **Achieve faster periodic convergence in case of cyclic loading** The computational expense of three-dimensional cardiovascular fluid dynamics simulations is often attributed to the need for computing multiple cycles (cardiac, respiratory, *etc*) before achieving a periodic solution. The work of [78] showed the possibility of fastening the periodic convergence by generating appropriate initial conditions using the simulation results of reduced-order 1D models.
- **Automating a comprehensive verification of CFD simulation quality** A few decades ago, multiple fluid mechanics editorial policy boards stated that “there is a need for higher standards on the control of numerical accuracy” and that “a single calculation of a fixed grid is not acceptable” [79]. However, recent research indicates that even grid refinement is insufficient for evaluating simulation quality. Instead, a comprehensive CFD investigation involving solver numerics, mesh and time-step refinement is essential. For instance, it has been shown in [80] that robust and minimally dissipative CFD solvers can tolerate surprisingly coarse resolutions, whereas solvers using low-order and/or stabilisation schemes may require much higher resolutions to detect relevant flow patterns.
- **Include Lumped Parameter Network (LPN) models for boundary conditions** While ensuring an accurate geometric representation of the computational domain is paramount in simulating fluid flow, it is equally essential to emphasise the importance of realistic boundary conditions. In general, the distribution of flow and pressure field within the simulated domain is often unknown and challenging to specify at the inflow/outflow boundaries. In fact, the physiology of the flow at the inlet/outlet is often too complex to be represented by a fixed-value Dirichlet of Neumann-type boundary condition. An alternative strategy involves linking the solution at the inflow/outflow boundaries of the modeled domain with simplified lumped parameter (0D models) or one-dimensional models representing the missing domain, check for instance [81][82]. This often requires solving an ODE to describe the physics of the flow at the boundary condition level.
- **Implementing Two-way coupling for embedded systems** Regarding the embedded systems, it is necessary to establish a two-way coupling between the embedded and the embedding domain. This coupling is necessary to ensure the conservation of mass (e.g. for mass transfer problems in vascular/tissue, root-soil and well-soil systems), momentum (e.g. to account for mechanical deformations in expanding or contracting airway/lung systems) and energy (e.g. for heat transfer problem in heat exchangers).

## 5. Conclusion

In this paper, we developed Tube2FEM, an open-source highly-automated framework for flow and flow-related processes simulation in (embedded) tubular structures. This system is capable of handling various input forms, including segmented tomographies and synthetic or skeletonised networks. It offers a range of features that include creating surface models, automated slicing and labeling for the assignment of boundary conditions. Furthermore, it supports surface and volumetric mesh generation and allows performing simulations using the Finite Element Method. It also offers capabilities for post-processing and visualising the results. The framework utilises a selection of carefully curated open-source softwares and libraries, chosen to guarantee a high degree of versatility for various simulation scenarios. Among the most significant of these are *GIBBON*, which is employed for image and/or geometry processing, *FEniCS* for conducting Finite Element (FE) simulations, and *Paraview*, which is utilised for the post-processing tasks.

The case studies presented in this work demonstrate the comprehensive capabilities of the Tube2FEM framework across a range of applications of (embedded) tubular geometries, from lung airways and root-soil interactions to microcirculation and tissue perfusion. We showcase in all case studies advanced meshing capabilities including surface mesh creation and extraction, isotropic remeshing, smoothing, anisotropy, multi-domains, and labelling for automated boundary conditions. Additionally, we showcase in case studies 4 and 5 a range of physics-based models applied to tubular and embedded tubular systems. Particularly, a framework for 3D-1D mixed dimensions simulation for embedded tubular system was created and was applied to simulate the solute transport from a microcirculation network to a surrounding biological tissue. In addition, a CT-to-simulation framework was also developed to conduct CFD and advection-diffusion-reaction simulations on image-based geometries.

Finally, future work will focus on enhancing the automation and efficiency of the workflow, expanding the framework’s capability to handle complex boundary conditions and couplings, and on integrating high performance solvers to achieve faster computational time while maintaining precision.

## Acknowledgments

The authors are grateful for funding from CRUK (C44767/A29458; HCS, SWS, RJS) and from the EPSRC (EP/W007096/1; HCS, SWS, RJS and EP/V034537/1; DO). The liver data of case study 5 were collected as part of an NIHR Invention for Innovation (i4i) Programme grant [II-LA-1116-20005] awarded to Prof Clarkson and Prof Davidson. Views expressed are those of the authors and not necessarily those of the NIHR or the Department of Health and Social Care.

## 6. Ethics Statement

**Lung Data:** Approval for this study of clinical and CT data was obtained from the local research ethics committees and Leeds East Research Ethics Committee: 20/YH/0120.

**Liver Data:** All data collected was with patient consent. The ethics approval for anonymized laparoscopic data collection was titled “Image guidance with image overlay for laparoscopic surgery” and was approved by the NRES Committee London – Stanmore. Reference No: 14/LO/1264 on 16/11/2017.

